# A Structurally Defined C12-Alkyl Quaternary Ammonium Compound reduces bacterial burden and biofilm in an infected wound model

**DOI:** 10.64898/2026.07.17.739236

**Authors:** Madelyn M. Kist, I Gusti Ayu I. Wulandari, Joseph F. Buell, Lisa A. Morici, Chad J. Roy

## Abstract

**Background:** Methicillin-resistant *Staphylococcus aureus* (MRSA) remains a common cause of skin and soft tissue infections, and topical agents that combine antimicrobial activity with wound compatibility are needed. C12-alkyl(ethylbenzyl)dimethyl ammonium chloride is a novel C12 quaternary ammonium molecule (hereafter, ‘EQ12’) related to benzalkonium chloride but designed to avoid the compositional heterogeneity and varied potency of conventional C8-C18 benzalkonium mixtures. We evaluated whether topical EQ12 is tolerated in healing wounds and whether it reduces MRSA burden in a splinted murine excisional wound infection model.

**Methods:** Female CD1 mice underwent 5-mm full-thickness dorsal excisional wounding, silicone splinting, and transparent dressing placement. For tolerability studies, uninfected wounds received EQ12 at 0.1 or 1 mg/mL or phosphate-buffered saline (PBS) and were followed for 14 days. For efficacy studies, wounds were inoculated with 1 × 10^4^ CFU MRSA and treated topically every 8 hours for 3 days beginning 4 hours after infection with EQ12 (1 mg/mL), bacitracin (500 U/mL), or PBS. Wound tissues were harvested on days 2, 4, and 6 for quantitative culture. Dressing-associated biofilm was evaluated by scanning electron microscopy.

**Results:** EQ12 did not alter body weight, wound inflammation scores, wound area, or percent wound closure compared with PBS in uninfected animals. In infected wounds, day 2 tissue burdens were not significantly different among treatment groups. By day 4, EQ12 significantly reduced MRSA burden compared with PBS, whereas bacitracin did not. By day 6, EQ12-treated animals had significantly lower MRSA burdens than both bacitracin- and PBS-treated controls. Mean tissue burden in the EQ12 group declined from 6.27 log10 CFU/g on day 4 to 5.53 log10 CFU/g on day 6, while bacitracin- and PBS-treated wounds remained persistently colonized. SEM revealed minimal adherent cocci or matrix-like bridging on dressings from EQ12-treated wounds, in contrast to dense microcolonies and biofilm-like structures on bacitracin and PBS dressings.

**Conclusions:** Topical EQ12 was compatible with gross wound healing and reduced MRSA burden in a dressed, splinted murine wound model. These data support continued development of EQ12 as a topical anti-staphylococcal wound-directed antimicrobial.

## Introduction

Skin and soft tissue infections (SSTIs) remain a frequent reason for outpatient evaluation, emergency department presentation, and hospital admission. Although SSTIs are clinically heterogeneous, wounds that breach the epidermal barrier are especially vulnerable to bacterial colonization and infection, and infected wounds can become self-sustaining microenvironments in which inflammation, tissue injury, impaired closure, and microbial persistence reinforce one another. *Staphylococcus aureus* is the dominant bacterial pathogen in culture-confirmed SSTIs, and methicillin-resistant *S. aureus* (MRSA) continues to complicate management because empiric therapy must account for resistance, local epidemiology, abscess formation, and patient comorbidity. For superficial or localized lesions, topical treatment is attractive because it can achieve high local exposure while limiting systemic toxicity and microbiome disruption. However, common topical antibiotics and antiseptics have important limitations, including resistance, cytotoxicity, narrow indications, staining, pain, and variable activity in the wound milieu. [1–4]

Topical bacitracin and mupirocin are widely used against Gram-positive skin pathogens, but both are constrained by resistance and by formulation-specific clinical indications. Classical antiseptics such as povidone-iodine, chlorhexidine, alcohols, hydrogen peroxide, and quaternary ammonium compounds (QACs) can rapidly reduce microbial burden on intact or prepared skin, but many are used primarily for prophylaxis or decontamination rather than repeated treatment of an open infected wound. A wound-directed antimicrobial intended for clinical translation must therefore satisfy two requirements simultaneously: it must reduce bacterial burden in the complex environment of injured skin, and it must avoid delaying the healing process it is intended to support. [5,6,21–23]

Biofilm biology further complicates the use of topical therapy in infected wounds. *S. aureus* can adhere to tissue and dressing surfaces, form microcolonies, and produce extracellular matrix that decreases antimicrobial penetration and increases phenotypic tolerance. In practice, this means that agents with excellent planktonic activity may not perform equivalently once bacteria are embedded in wound exudate, necrotic debris, host proteins, and polymeric dressing interfaces. Topically administered therapies should therefore emphasize not only bacterial killing but also whether treatment alters the local wound ecology that enables persistence. The present study addresses this translational gap by pairing quantitative tissue culture with SEM of dressing-associated bacterial architecture, while reserving the detailed *in vitro* antibiofilm for a separate mechanistic manuscript reported elsewhere. [5,12,24]

C12-alkyl(ethylbenzyl)dimethyl ammonium chloride (C23H42ClN), or otherwise ‘EQ12’, is a structurally defined alkylbenzyl dimethyl ammonium compound. The molecule is conceptually related to benzalkonium chloride (BZK), which is typically a mixture of alkyl-chain homologs, but EQ12 is represented as a single C12-alkyl configuration. This is a central chemical distinction. QAC activity depends on the cationic head group and hydrophobic alkyl chain: electrostatic interaction with negatively charged bacterial surfaces is followed by membrane insertion, disruption of membrane integrity, and cell death. Alkyl-chain length influences potency, membrane insertion, and cytotoxicity, making a defined C12 species a plausible strategy to tune antimicrobial effect while limiting host-cell damage. [6–9]

The *in vitro* antibacterial properties of C12-QAC are being reported separately and are not reproduced here in detail. Briefly, those studies support the use of a 1 mg/mL topical working concentration for animal testing: the MRSA minimum inhibitory and bactericidal concentrations were both 4.7 µg/mL, making the *in vivo* working solution approximately 213-fold above the MRSA MIC. In separate fibroblast-based cytotoxicity testing, effective concentrations (1 mg/ml) of EQ12 graded a categorical ‘0’, or as normal saline (data not shown). These supportive observations provided a rationale for testing topical EQ12 in a live wound model, but the present manuscript focuses on *in vivo* tolerability, MRSA tissue burden, and dressing-associated biofilm morphology. [24,25]

We used a splinted full-thickness murine excisional wound model with transparent adhesive dressing. This model was selected because unsplinted rodent wounds close primarily by contraction, whereas splinting forces a more clinically relevant healing pattern involving re-epithelialization and granulation. The addition of a transparent dressing retains inoculum and treatment within the wound bed and approximates aspects of contemporary wound care. We hypothesized that topical EQ12 would be grossly compatible with wound healing and would reduce MRSA burden relative to PBS vehicle and bacitracin comparator treatment. [10–13]

## Methods

### Study agent and comparators

EQ12 was supplied as C12-alkyl(ethylbenzyl)dimethyl ammonium chloride and prepared in sterile phosphate-buffered saline (PBS) for topical administration. The 1 mg/mL working concentration was selected based on prior *in vitro* MRSA susceptibility and cytotoxicity data. A lower 0.1 mg/mL concentration was included in the uninfected wound-healing study to assess dose-related tolerability. Bacitracin (500 U/mL) served as a clinically familiar topical antibiotic comparator for MRSA-infected wounds. Sterile 1X PBS served as vehicle and negative control. [6–9]

### Animals and ethics

All animal procedures were performed in accordance with the Guide for the Care and Use of Laboratory Animals and approved by the Tulane University Institutional Animal Care and Use Committee under protocol 2162. Female CD1 mice, 8 to 10 weeks old, were obtained from Charles River Laboratories. Animals were maintained under specific pathogen-free conditions with alfalfa-free rodent chow and acidified water *ad libitum*. Before surgery, animals were group housed in standard microisolator cages; after wounding, animals were individually housed to minimize suture disruption, dressing manipulation, and transfer of inoculum or topical treatment between animals. [17]

### Splinted full-thickness excisional wound model

Animals received ketamine/xylazine anesthesia (90 mg/kg and 10 mg/kg, respectively) diluted in PBS and administered intraperitoneally. Adequate anesthesia was confirmed by footpad reflex. Extended-release buprenorphine (3.25 mg/kg) was administered subcutaneously before surgery for analgesia, and ophthalmic ointment was applied. Dorsal fur was removed from the base of the skull to approximately 3 cm caudally and between the scapulae. The operative field was disinfected with sequential passes of 2% chlorhexidine gluconate and 70% ethanol. A sterile 5-mm biopsy punch was used to outline a circular wound in the dorsal region near the base of the neck, aligned with the scapulae. The tissue was elevated with forceps and excised with sterile scissors to create a full-thickness defect extending through skin into the subcutaneous layer. To limit contraction, a silicone torus generated from medical-grade silicone was secured around the wound with cyanoacrylate and eight interrupted 4-0 silk sutures; knots were reinforced with cyanoacrylate. Wounds were covered with pre-cut circular Tegaderm transparent dressing. Animals recovered on a temperature-controlled heating pad and were returned to individual cages when ambulatory. [10–13]

### Uninfected wound-healing study

To evaluate tolerability in the absence of infection, uninfected wounds were treated topically with EQ12 at 0.1 mg/mL, C12-QAC at 1 mg/mL, or PBS. Treatments were administered through the dressing by piercing the Tegaderm with an insulin syringe and applying 20 µl directly to the wound bed. Animals were monitored through day 14. Body weight was recorded daily. Wounds were scored categorically as 0, no redness, swelling, or exudate; 1, light inflammation or some exudate; 2, redness, swelling, exudate, or discoloration; or 3, heavy inflammation, discoloration, and exudate or purulence. Beginning on day 4 after dressing removal, wound diameters were measured daily with a ruler. Wound area was calculated as *pi*\*r^2^, and percent closure was calculated relative to the original 5-mm wound area.

### MRSA wound infection and topical treatment

For efficacy studies, bacterial cultures were grown and harvested at an optical density at 600 nm of 0.75 for MRSA. Cells were resuspended in PBS and diluted to 1 × 10^6^ CFU/mL. Using a 0.5-mL insulin syringe to pierce the Tegaderm dressing, each wound received 10 µl of inoculum, delivering 1 × 10^4^ CFU directly into the open wound bed. Beginning 4 hours after surgery and infection, mice were briefly anesthetized with inhaled isoflurane and treated topically with 20 µl of EQ12 (1 mg/mL), bacitracin (500 U/mL), or PBS. Treatments were administered every 8 hours, within a 1-hour window, through day 3 for a total of nine treatments. Animals were assigned to staggered harvest groups with three animals per treatment group per time point, except where noted. [12,13]

### Quantification of tissue bacterial burden

Animals were euthanized on days 2, 4, or 6 after infection by carbon dioxide asphyxiation followed by cervical dislocation. Splints, sutures, and dressings were removed aseptically. Wound tissue was excised as a box around the biopsy site, including epidermis, dermis, subcutaneous tissue, and underlying smooth muscle. Tissue was weighed, placed in PBS, minced, and homogenized. Because fur and smooth muscle increased sample viscosity, a modified dilution series was used before plating. Homogenates were plated on mannitol salt agar for MRSA selection and incubated at 37°C for 24 hours. Colonies were enumerated and normalized to tissue mass as CFU/g tissue. Data are reported as mean log10 CFU/g tissue. The day 6 PBS group included two animals because one animal was excluded before surgery because of preoperative complications.

### Scanning electron microscopy of wound dressings

On day 4 after infection, Tegaderm dressings from representative animals were removed and processed for scanning electron microscopy (SEM). A section directly overlying the wound bed was excised, mounted on a hydroxyapatite disc using cyanoacrylate, air dried, and fixed in 2% electron microscopy-grade glutaraldehyde in 0.1 M phosphate buffer at 4°C for 16 to 18 hours. Samples were washed in water, dehydrated through graded ethanol, critical-point dried, mounted with conductive silver tape, carbon coated, and imaged with a Hitachi 3400 scanning electron microscope at 2,000x and 10,000x magnification. [24]

### Statistical analysis

Analyses and figures were generated using GraphPad Prism 10.3.1. Body weight, wound scores, wound area, wound closure, and bacterial burdens were compared across treatment groups using ordinary one-way analysis of variance with multiple comparisons, as appropriate for each time point or outcome. Statistical significance was defined as *p*<0.05.

## Results

### Topical EQ12 did not impair gross wound healing in uninfected animals

Across the 14-day uninfected wound-healing study, topical EQ12 produced no overt evidence of impaired healing or local intolerance. Body weight trajectories were comparable among animals treated with 0.1 mg/mL EQ12, 1 mg/mL EQ12, or PBS. Wound scores also remained low and similar across groups, without treatment-associated increases in redness, swelling, discoloration, exudate, or purulence. Quantitative wound measurements supported the clinical observations. Wound areas declined over time in all groups, and percent closure curves were overlapping. By the end of follow-up, wound closure approached completion regardless of treatment. Together, these data indicate that repeated topical EQ12 exposure at the working concentration used for efficacy testing did not measurably interfere with gross healing dynamics in this model.

**Figure 1.**
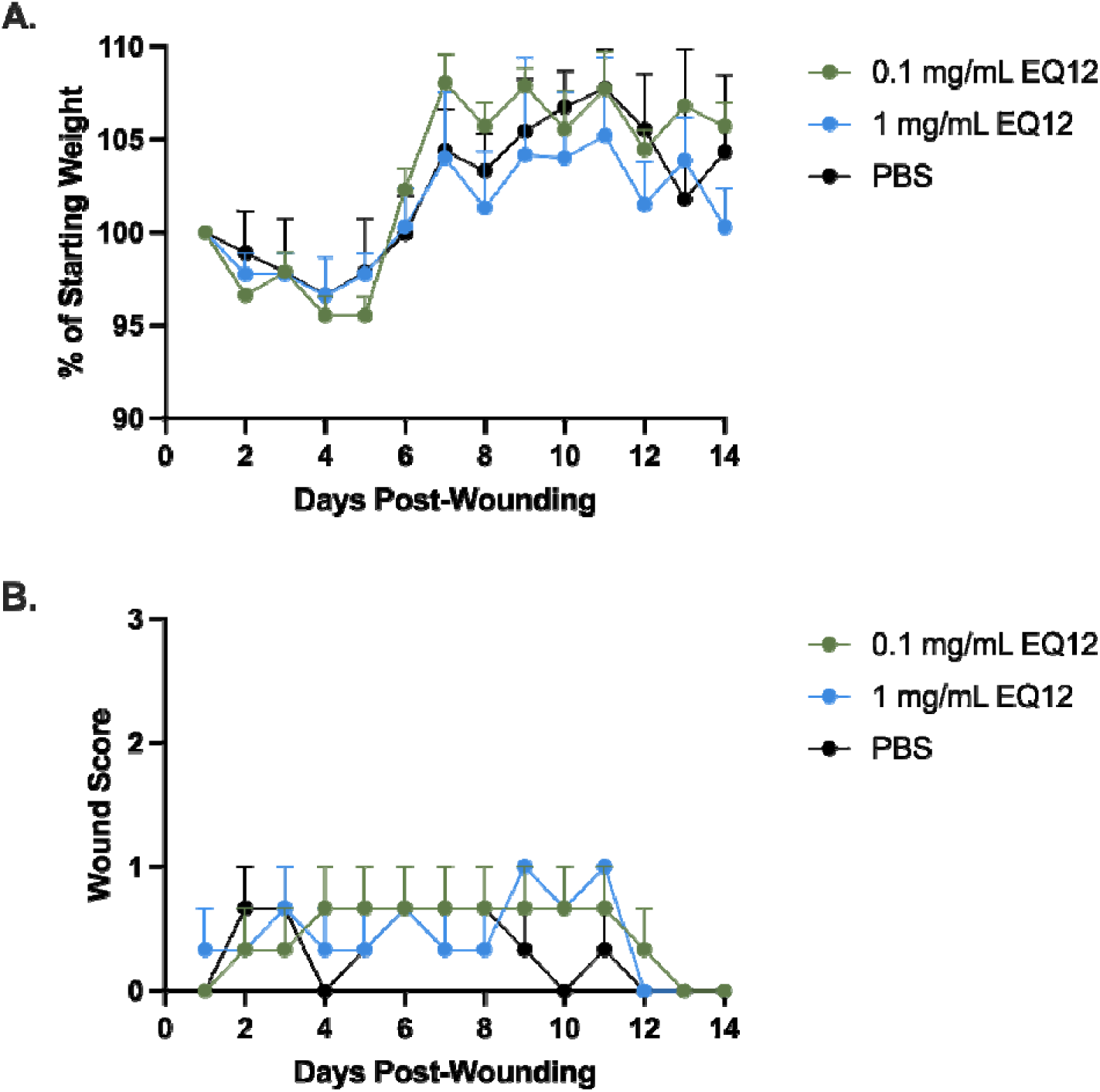
EQ12 does not affect body weight or wound scores in uninfected mice. Animals were monitored, weighed, and scored daily from day 0 through day 14. Wounds were scored as 0 to 3 based on inflammation, exudate, discoloration, and purulence. Data represent biological replicates plotted as mean +/− SEM. No significant differences were observed among groups.

**Figure 2.**
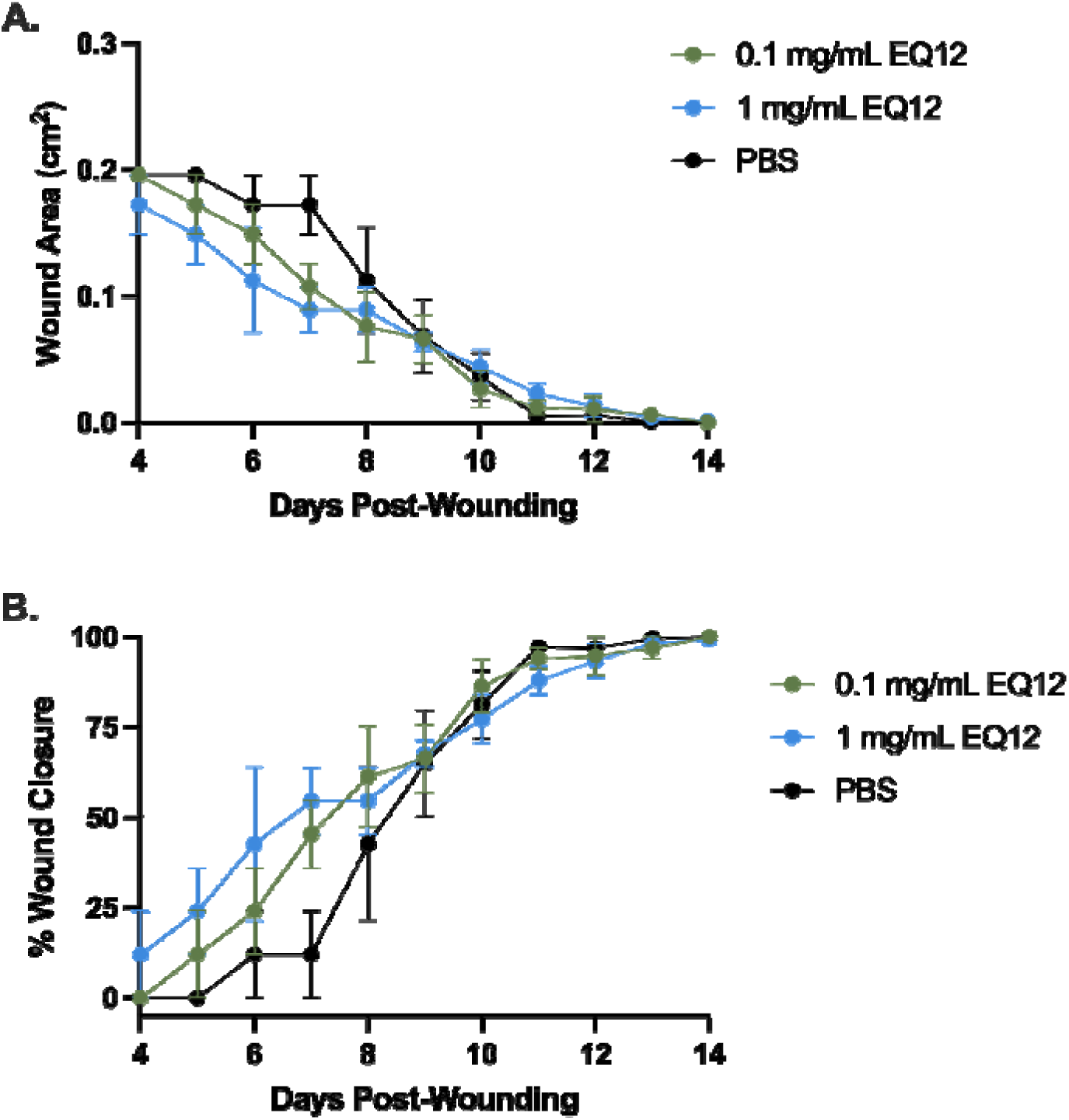
EQ12 does not impair wound closure in uninfected mice. Wound diameter was measured daily beginning on day 4 after removal of the transparent dressing through day 14. Wound area and percent closure were calculated relative to the 5-mm initial wound. Data represent biological replicates plotted as mean +/− SEM. No significant differences were observed among groups.

### EQ12 reduced MRSA tissue burden after repeated topical treatment

In MRSA-infected wounds, tissue burdens on day 2 did not differ significantly among EQ12, bacitracin, and PBS groups, indicating that a separation in efficacy was not yet detectable early after infection and treatment initiation. By day 4, however, EQ12 significantly reduced MRSA burden compared with PBS, whereas bacitracin did not differ significantly from PBS. By day 6, the difference was more pronounced: EQ12-treated wounds had significantly lower MRSA burdens than both bacitracin-treated wounds and PBS controls, while bacitracin remained statistically indistinguishable from PBS. Time-course analysis provided additional context. Mean MRSA burden in EQ12-treated wounds increased from 5.89 log10 CFU/g on day 2 to 6.27 log10 CFU/g on day 4, then declined to 5.53 log10 CFU/g by day 6. In contrast, bacitracin-treated wounds remained persistently burdened, with mean values of 6.65, 6.49, and 6.97 log10 CFU/g on days 2, 4, and 6, respectively. PBS-treated wounds similarly remained high, with mean values of 7.58, 7.03, and 7.38 log10 CFU/g. These results suggest that EQ12 did not sterilize the wound bed under the tested regimen but did reduce MRSA proliferation and improve bacterial control relative to both untreated and bacitracin-treated wounds.

**Figure 3.**
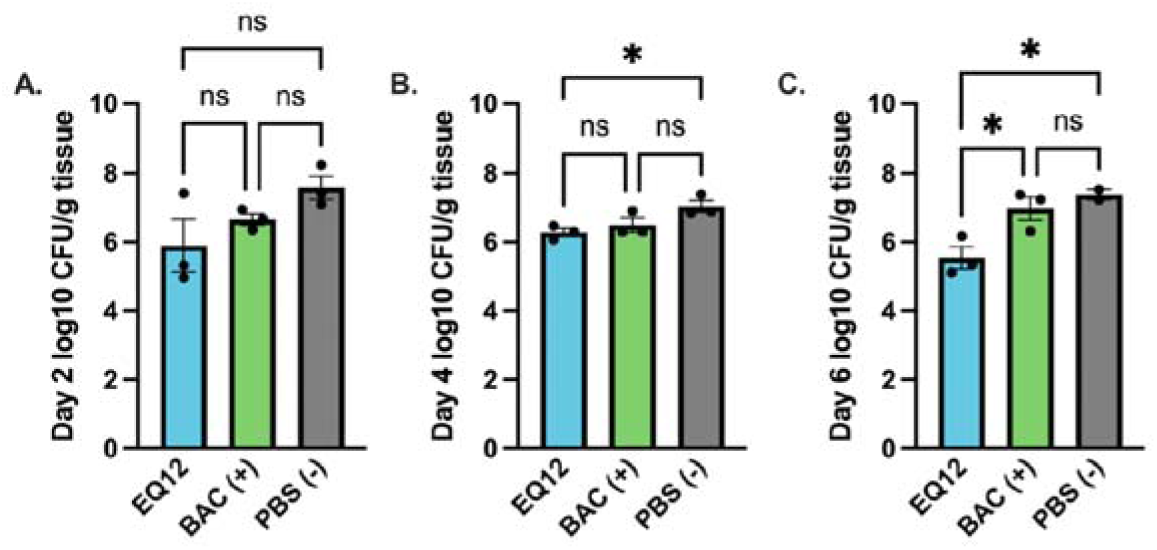
Topical EQ12 reduces MRSA burden in wound tissue on days 4 and 6. MRSA-infected wounds were treated every 8 hours from 4 hours after infection through day 3 with EQ12 (1 mg/mL), bacitracin (500 U/mL), or PBS. Wound tissues were harvested on days 2, 4, and 6 and plated on mannitol salt agar. Data are mean +/− SEM. \**p* < 0.05.

**Figure 4.**
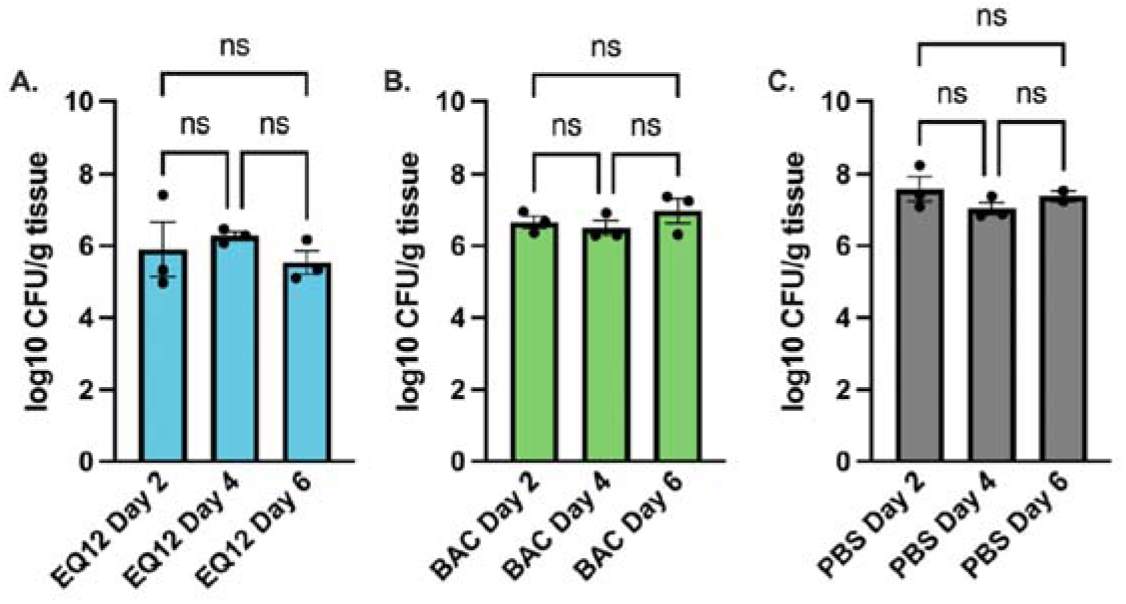
Time course of MRSA burden after topical C12-QAC, bacitracin, or PBS. MRSA burdens from Figure 3 are plotted by treatment group across time. Data are mean +/− SEM. No within-treatment time-course comparisons reached significance in the source analysis.

### EQ12 limited MRSA biofilm-like growth on wound dressings

SEM of Tegaderm dressings removed on day 4 provided qualitative evidence that topical EQ12 altered the dressing-associated bacterial architecture. Dressings from EQ12-treated wounds showed a smooth dressing surface without obvious adherent cocci, dense microcolonies, or matrix-like bridging consistent with staphylococcal biofilm. In contrast, dressings from bacitracin-treated animals contained visible clusters of cocci and microcolonies, and PBS control dressings showed extensive adherent bacterial aggregates, extracellular material, and mesh-like matrix bridging. The distinction was most apparent at higher magnification, where bacitracin and PBS samples contained dense staphylococcal packing while EQ12-treated samples lacked comparable surface-associated biofilm features. These findings support the quantitative culture results and suggest that the benefit of EQ12 may extend beyond planktonic bacterial killing to interference with wound-dressing-associated MRSA organization.

**Figure 5.**
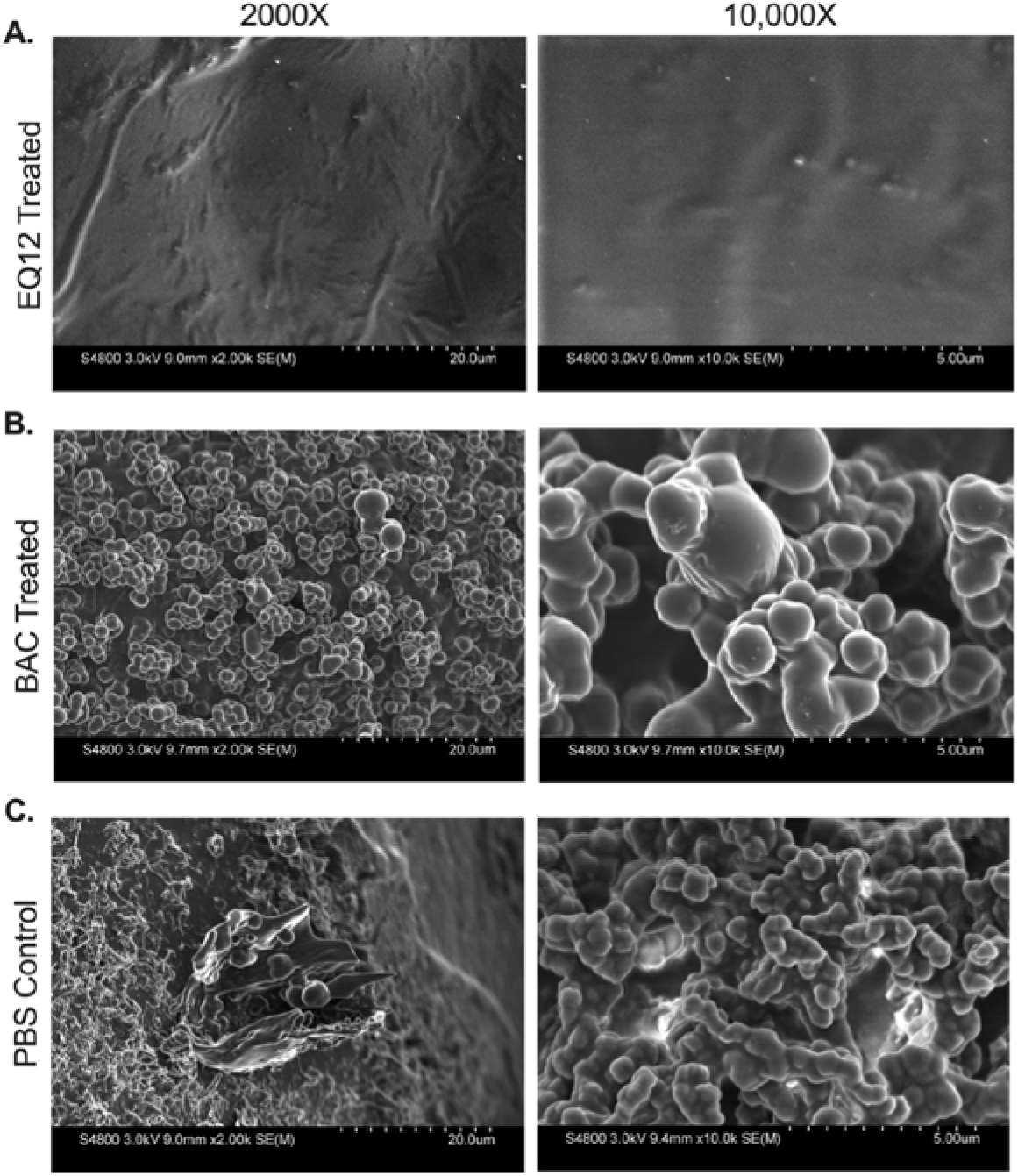
EQ12 limits MRSA biofilm-like growth on wound dressings. Tegaderm dressings were removed from representative infected wounds on day 4 and imaged by scanning electron microscopy at 2,000x and 10,000x. Dressings from EQ12-treated wounds lacked dense adherent cocci and matrix-like bridging compared with bacitracin and PBS controls.

## Discussion

This study demonstrates that topical EQ12 is compatible with gross wound healing and reduces MRSA burden in a dressed, splinted murine excisional wound infection model. The findings are clinically relevant for three reasons. First, MRSA remains a major cause of SSTI and wound-associated morbidity. Second, repeated topical therapy must be evaluated in an injured-skin context rather than inferred from in vitro susceptibility alone. Third, the treatment was compared with bacitracin, a familiar topical antibiotic comparator, and showed superior bacterial control under the tested conditions. [3–5,12,13]

The chemical identity of the investigational agent is central to interpretation. Conventional benzalkonium chloride preparations are mixtures of alkyl-chain homologs, whereas EQ12 is represented as C12-alkyl(ethylbenzyl)dimethyl ammonium chloride, a structurally defined C12 alkyl isomeric QAC. This distinction matters because QAC function depends on both charge-driven surface interaction and hydrophobic membrane insertion. A structurally defined C12-isomer molecule provides a more consistent balance between antimicrobial activity and host-cell compatibility than a heterogeneous homolog mixture. The present data do not define the molecular mechanism *in vivo*, but they support the translational premise that a chemically defined EQ12 can be administered repeatedly to open wounds without grossly delaying closure while still reducing MRSA burden. [6–9]

The splinted wound model strengthens the translational value of the results. Rodent wounds can close rapidly by contraction, which can confound studies of topical antimicrobials and wound repair. Silicone splinting limits contraction and forces closure through re-epithelialization and granulation, features more consistent with human healing. The Tegaderm dressing provides an additional clinically relevant surface where bacteria can adhere and organize. This aspect of the model is important because wound care often involves dressings, and a topical antimicrobial must function in the presence of tissue fluid, host debris, polymeric dressing surfaces, and repeated handling. [10–13,17,18]

The tolerability study is an important component of the manuscript because antimicrobial potency alone is insufficient for wound-directed therapy. Several topical antiseptics can be effective antimicrobials but may be unsuitable for repeated use in open wounds because of cytotoxicity or concerns for delayed healing. In uninfected wounds, repeated EQ12 treatment did not significantly alter body weight, inflammatory wound score, wound area, or closure rate compared with PBS. These gross outcomes do not exclude subtler histologic or molecular effects, but they provide an initial *in vivo* safety signal at the 1 mg/mL concentration selected for efficacy testing. [5,21,22]

The MRSA burden data suggest a delayed but meaningful treatment effect. The lack of a significant difference on day 2 may reflect the early establishment phase of infection, the limited number of doses administered by that time, and the complexity of the wound environment. By day 4 and day 6, after repeated dosing, EQ12 separated from PBS and bacitracin. This pattern is clinically relevant because topical wound therapies are generally administered over days rather than as a single decontamination event. Importantly, bacitracin did not significantly reduce tissue burden relative to PBS in this model, emphasizing the need to evaluate commonly used topical comparators in models that include wound architecture and dressing-associated bacterial persistence. [12,13,23]

From a therapeutic positioning standpoint, the most relevant comparator in this study is not an optimized systemic anti-MRSA regimen but a commonly used topical antibiotic. Bacitracin is familiar to clinicians and patients, yet its activity is limited to selected Gram-positive organisms and it is not designed to overcome the full biological complexity of an established wound infection. The finding that EQ12 separated from bacitracin by day 6 suggests that a chemically defined topical antiseptic-like compound may be useful when framed as local bacterial-burden control rather than as a conventional systemic antibiotic substitute. This distinction frames EQ12 for development as a wound-directed topical antimicrobial that could complement debridement, dressing changes, source control, and systemic therapy when indicated, not a replacement for clinical management of invasive disease. [5,21–23]

The SEM findings are hypothesis-generating and important. Dressings from control and bacitracin-treated wounds showed coccal aggregates and extracellular material consistent with MRSA biofilm-like organization, whereas dressings from EQ12-treated wounds lacked comparable structures. Biofilm formation on dressings and wound surfaces can contribute to persistence, tolerance, and recurrent inflammation. The current SEM analysis was qualitative and performed on representative samples, so it should not be interpreted as a quantitative antibiofilm endpoint. Nonetheless, the imaging supports the conclusion that EQ12 changes the wound-dressing microbial phenotype in a manner consistent with reduced surface-associated MRSA organization. [24]

This work has limitations. The sample size was modest, with three animals per treatment group per time point for infected studies and one day 6 PBS exclusion. Only female CD1 mice were used. Sex, strain, age, and immune status can influence wound healing and *S. aureus* pathogenesis, and future studies should evaluate whether efficacy is maintained across more diverse host backgrounds. The study relied on gross wound-healing endpoints and quantitative culture rather than histopathology, cytokine profiling, tissue penetration, or pharmacokinetic/pharmacodynamic measurements. The topical dosing regimen was also limited to every 8 hours through day 3. Longer treatment, alternative vehicles such as ointment or cream bases, and concentration-response studies could further define the therapeutic window. Finally, although supportive *in vitro* data informed dose selection, this manuscript intentionally does not reproduce the broader *in vitro* dataset as those studies are being reported separately. [17–20,25,26]

Several development questions are especially important before clinical translation. The current liquid formulation ensured standardized dosing through the transparent dressing, but cream, gel, hydrogel, or impregnated-dressing formats may increase residence time and patient usability. Pharmacokinetic and pharmacodynamic studies should define how much active compound remains in the wound bed, how rapidly it partitions into tissue and dressing material, and whether repeated administration alters local inflammation or epithelial migration. Histology could determine whether EQ12 changes granulation tissue, angiogenesis, collagen deposition, or re-epithelialization despite normal gross closure. Finally, testing against multiple clinical MRSA isolates, including mupirocin-resistant isolates, would strengthen the medical relevance of the anti-staphylococcal claim. [18,22,23,25,26]

In conclusion, topical EQ12 reduced MRSA wound burden and limited dressing-associated biofilm-like growth without measurably impairing gross healing in a murine splinted excisional wound model. These findings support continued preclinical development of EQ12 as a chemically defined topical anti-staphylococcal agent for wound-associated SSTI. Future work should prioritize dose optimization, formulation development, histologic assessment, pharmacokinetic profiling, and confirmation in larger or more clinically analogous wound models. [12,13,26]

## Declarations

## Ethics approval

All animal procedures were approved by the Tulane University Institutional Animal Care and Use Committee under protocol 2162 and conducted in accordance with the Guide for the Care and Use of Laboratory Animals.

## Data availability

Data underlying this draft are derived from the uploaded dissertation and poster source materials. Raw data files should be deposited or made available according to the target journal policy before submission.

## Acknowledgments

The authors thank the Tulane animal care, imaging, and laboratory support teams.

